# The genomic signature of wild-to-crop introgression during the domestication of scarlet runner bean (*Phaseolus coccineus* L.)

**DOI:** 10.1101/2021.02.03.429668

**Authors:** Azalea Guerra-García, Idalia C. Rojas-Barrera, Jeffrey Ross-Ibarra, Roberto Papa, Daniel Piñero

**Affiliations:** Departamento de Ecología Evolutiva, Instituto de Ecología, Universidad Nacional Autónoma de México, Ciudad de México, México; Department of Plant Sciences, University of Saskatchewan, Canada; Environmental Genomics, Max Planck Institute for Evolutionary Biology, 24306 Plön, Germany; Department of Evolution and Ecology, Center for Population Biology, and Genome Center, University of California, Davis, CA, USA; Dipartimento di Scienze Agrarie, Alimentari ed Ambientali, Università Politecnica delle Marche, 60131 Ancona, Italy

## Abstract

- The scarlet runner bean is an open-pollinated legume from the highlands of Mesoamerica that is cultivated in small-scale agriculture for its dry seeds and immature pods. Demographic bottlenecks associated with domestication might reduce genetic diversity and facilitate the accumulation of deleterious mutations. Conversely, introgression from wild relatives could be a source of variation.
- Using Genotyping by Sequencing data (79,286 SNVs) from 237 cultivated and wild samples, we evaluated the demographic history of traditional varieties from different regions of Mexico and looked for evidence of introgression between sympatric wild and cultivated populations.
- Traditional varieties have high levels of diversity, even though there is evidence of a severe initial genetic bottleneck, followed by a population expansion. Introgression from wild to domesticated populations was detected, but not in the opposite direction. This asymmetric introgression might contribute to the recovery of genetic variation and it has occurred at different times: constantly in the center of Mexico; recently in the North West; and anciently in the South.
- Several factors are acting together to increase and maintain genetic diversity in *P. coccineus* cultivars, such as demographic expansion and introgression. Wild relatives represent a valuable genetic resource and have played a key role in scarlet runner bean evolution via introgression into traditional varieties.

## Introduction

The scarlet runner bean (*Phaseolus coccineus* L.) is one of the five domesticated *Phaseolus* species. It is a close relative of common bean (*P. vulgaris*) and year bean (*P. dumosus*), which were also domesticated in Mesoamerica. In contrast with common bean that is predominantly autogamous, year bean and scarlet runner bean are allogamous species (Bitocchi *et al*., 2017). Scarlet runner bean is a perennial species, inhabiting the highlands of Mesoamerica (1,000-3,000 m.a.s.l), from northern Mexico (Chihuahua) to Panama, although it is usually cultivated as an annual crop for dry seed and immature pods (Delgado-Salinas, 1988).

Because of the high phenotypic variation of runner bean, Freytag & Debouck (2002) proposed two subspecies: *P. coccineus* subsp. *coccineus*, a red-flowered type including 11 varieties (one of these is the cultivated form); and *P. coccineus* var. *striatus*, a purple-flowered type with eight wild varieties.

The domesticated form of *P. coccineus* is cultivated in Mexico, Guatemala and Honduras (Delgado-Salinas, 1988), and due to its tolerance to cold, it is also cultivated in European countries such as the United Kingdom, Netherlands, Italy and Spain (Rodiño *et al*., 2007).

Few studies have focused on the domestication history of scarlet runner beans. Initially, two domestication events were suggested using low-resolution molecular markers (Spataro *et al*., 2011; Rodriguez *et al*., 2013). Nevertheless, those works have mostly focused on European cultivars. More recently, (Guerra-García *et al*., 2017) proposed just one domestication of *P. coccineus*, which probably took place in the central Mexican biogeographic area known as the Trans Mexican Volcanic Belt (TMVB).

The demographic history of crops shapes patterns and levels of genetic variation on which natural and artificial selection can act (Gaut *et al*., 2018). The first stages of domestication are often associated with genetic bottlenecks because early farmers likely used a limited number of wild individuals for initial management and cultivation (Meyer & Purugganan, 2013; Gaut *et al*., 2018). Domestication is a gradual process in which both population size changes and gene flow between the wild relatives and the incipient crops play a role in determining overall levels of genetic variation. The subsequent range expansion out of the center of origin leads to the adaptation of the domesticated species to different environments as well as distinct cultural preferences (Meyer & Purugganan, 2013; Gaut *et al*., 2015; Janzen *et al*., 2019). The role of hybridization in domestication has been widely documented (Stewart *et al*., 2003; Arnold, 2004; Hancock, 2012; Bredeson *et al*., 2016; Choi & Purugganan, 2018) and evidence suggests that wild-to-crop introgression and even interspecific hybridization, can be a source of crop adaptation and may play a role in crop diversification and range expansion after domestication (Janzen *et al*., 2019; Purugganan, 2019). One of the most relevant examples of adaptive introgression in crops is the case of maize adaptation to highlands as a result of introgression from wild populations of *Zea mays* ssp. *mexicana* (van Heerwaarden *et al*., 2011; Hufford *et al*., 2013; Takuno *et al*., 2015).

Demographic bottlenecks associated with domestication might lead to a shift in the effectiveness of selection. Because population size is reduced during bottlenecks, selection is expected to be less efficacious in removing deleterious mutations (Morrell *et al*., 2012; Moyers *et al*., 2018). In domesticated forms the increased genetic load is called the ‘cost of domestication’ and it has been documented in species like rice (Lu *et al*., 2006), maize (Mezmouk & Ross-Ibarra, 2014), sunflower (Renaut & Rieseberg, 2015), and cassava (Ramu *et al*., 2017). In this work, we provide an analysis of the population structure of wild and domesticated populations of scarlet runner bean and investigate its demographic history during its domestication and subsequent spread, the role of gene flow and introgression between wild and domesticated populations, and how these processes have shaped the genetic diversity present in the cultivars and wild populations of *P. coccineus* in Mexico.

## Materials and Methods

### Sampling and genomic data

Plant material was collected from Northwest (Durango) to Southeast of Mexico (Chiapas) during 2014 and 2015. Wild individuals were sampled in nine locations, ferals in two sites and traditional varieties in 11 geographic points. Samples of the breeding line Blanco Tlaxcala and a cultivar from Spain were also included (Table S1). Categories (wild, feral, traditional variety) were assigned according to habitat and morphological observations. Only one of the wild populations corresponded to subsp. *striatus*. Samples from the closely related species *P. vulgaris* and *P. dumosus* were additionally included and used as outgroups.

Leaf tissue from wild samples was collected and stored in silica until processed. Seeds from traditional varieties were germinated at the Instituto de Ecología, UNAM. DNA was extracted using a DNeasy Plant Mini Kit (Qiagen). Library preparation and sequencing were performed at the Institute for Genomic Diversity at Cornell University. For library construction, double digestion was performed using PstI and BfaI enzymes, following Elshire *et al*. (2011). A total of 326 samples were sequenced in four lanes of an Illumina HiSeq 2500 (100 bp, single-end reads).

### Variant discovery, genotyping and filtering

Fastq files were demultiplexed with GBSx 1.3 (Herten *et al*., 2015) and reads were trimmed with Trimmomatic 0.36 (Bolger *et al*., 2014). Alignments were performed with Nextgenmap 0.5.3 (Sedlazeck *et al*., 2013) using the *Phaseolus vulgaris* genome v2.1 (DOE-JGI and USDA-NIFA, http://phytozome.jgi.doe.gov/) and then were converted to binary files using samtools 1.5 (Kaisers *et al*., 2015). Single Nucleotide Variants (SNVs) were discovered for each sample using the HaplotypeCaller tool and genotypes were then merged with GenotypeGVCFs. Both tools are from the Genome Analysis Toolkit (GATK 4.0.1.0; (McKenna *et al*., 2010).

VCFtools 0.1.15 (Danecek *et al*., 2011) was used to perform the variant filtering according to the following parameters: minimum mean depth 6X; max missingness per sample 0.30; max missingness per site 0.05; loci not mapped in *P. vulgaris* chromosomes were excluded; and only biallelic sites were kept. Also, SNVs that were not in Hardy-Weinberg equilibrium (*p* < 0.01) in at least one wild population were identified with PLINK 1.07 (Chang *et al*., 2015) and filtered, as well as the 15,601 putative paralogs detected with HDplot (McKinney *et al*., 2017).

We classified the SNVs into three categories: non-genic, intronic and coding regions (CDS). The consequence of SNVs within coding regions was predicted with the R package VariantAnnotation (Obenchain *et al*., 2014).

### Defining “populations”

Diversity analyses were performed at the “population” level. Populations were established according to 1) Principal Component Analysis (PCA) performed with SNPrelate (Zheng *et al*., 2012); 2) the genetic groups identified with Admixture v1.3 (Alexander *et al*., 2009); and 3) the topology of the phylogenetic hypothesis constructed with FastTree (Price *et al*., 2009). Populations may differ from locations because in some cases individuals from different locations did not show differentiation and as a consequence belonged to the same genetic group. In other cases, genetic groups were split because a clear differentiation was observed in the PCA and in the phylogenetic tree. The nature of the samples was also considered (e.g. feral, breeding line or traditional variety).

### Measuring diversity

Heterozygosity and inbreeding coefficient (*F*_IS_) per site were estimated with Hierfstat package (Goudet, 2005) according to the established populations, performing a bootstrap (1,000) to obtain confidence intervals for the inbreeding coefficient, and Kruskal-Wallis and Pairwise test to compare the heterozygosity among populations. A custom R script was made to discover the private SNVs within each population, considering only the polymorphic sites within the groups. This R script uses the Hierfstat package (Goudet, 2005) to estimate allele frequencies. It is important to note that the amount of polymorphic sites and private alleles within each population is dependent on sample size (different amount of individuals was available per population) and as a consequence, we applied a rarefaction approach for allelic richness and private allelic richness using ADZE v1.0 (Szpiech *et al*., 2008), excluding loci with missing data greater than 0.2 for at least one population.

We tested the hypothesis that genetic diversity of landraces decreases with increasing distance from the center of domestication. For this, a correlation was performed using heterozygosity and distance from the centroid of the TMVB traditional varieties to the rest of the cultivated populations. Breeding line Blanco Tlaxcala and cultivar from Spain were not included in this analysis.

### Detecting introgression and gene flow

Two approaches were used to infer gene flow: TreeMix (Pickrell & Pritchard, 2012) and Patterson’s D, also known as the ABBA-BABA test (Green *et al*., 2010; Durand *et al*., 2011). The gene flow scenarios obtained with TreeMix were then tested with the ABBA-BABA approach, as well as gene flow among sympatric populations.

The ABBA-BABA test is based on a resolved phylogeny among four taxa (((H1, H2), H3), H4) and determines if the pattern of derived alleles is consistent with the phylogeny (Green *et al*., 2010; Durand *et al*., 2011). To compute this test we used bam files from each individual, and ran the analysis with the multipop ABBA BABA module from the package ANGSD (Korneliussen *et al*., 2014). A positive D value indicates gene flow from H3 into H2 (ABBA) and a negative value indicates gene flow from H3 into H1 (BABA).

Gene flow events obtained from TreeMix (see results) that were evaluated with the ABBA BABA test included those with: 1) old gene flow among the branch of traditional varieties from TMVB and SMOCC (Cult-TMVB and Cult-SMOCC) and wild populations from TMVB (Wild-TMVB); 2) an ancient admixture involving the branch of all cultivars and the wild population from Chiapas (Wild-SUR-CH); 3) between ferals and the wild population from Chiapas (Wild-SUR-CH). Scenarios of gene flow between sympatric wild populations and traditional varieties, and between the sympatric *P. dumosus* cultivars and Chiapas populations (Wild-SUR-CH and Cult-SUR) were also evaluated. For all tested scenarios, wild *P. vulgaris* was used as an outgroup (H4), and the statistical significance (*p* < 0.05) was established after applying a Bonferroni correction to the block jackknife p-value.

### Inferring the demographic history of scarlet runner bean

To find evidence of demographic processes that have affected *P. coccineus* populations, the Site Frequency Spectrum (SFS) of each population was constructed using the PLINK allele count function. Because the different SNV categories may be under different evolutionary processes, they provide complementary information. Therefore we constructed the SFSs according to SNV in CDS, non-genic and intronic regions. The expected SFS was derived by estimating 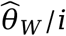, where i is the number of individuals per population. To identify signs of demographic bottlenecks, Long Runs of Homozygosity (ROH) per individual were estimated with PLINK using a 500 Kb min window size.

With the patterns observed in the SFSs, the ROHs and the introgression events found with TreeMix and the ABBA-BABA test, demographic scenarios were constructed and tested using fastsimcoal2. fastsimcoal2 uses coalescent simulations to model demographic scenarios from the SFS (Excoffier *et al*., 2013). Again, only non-genic regions were included to make the demographic inferences. For the Cult-TMVB, Cult-SMOCC and Cult-SUR-CH three scenarios were modeled. The models differ in when introgression from the wild relative occurred: recent (3,000 generations ago to present), ancient (from 6,000 to 3,000 generation ago), and constant (6,000 generation ago to present for Cult-SMOCC and Cult-SUR-CH, and from divergence time to present for Cult-TMVB). Additionally, bidirectional introgression was included for the TMVB populations during the first 2,000 generations after divergence, which we considered as an early domestication phase.

In these demographic models, the cultivated population diverged from the wild one TDIV (time of divergence) generations ago in the case of Cult-SMOCC and Cult-SUR-CH, and TDOM (time of domestication) generations ago for the Cult-TMVB. After an initial bottleneck in the cultivated population (NAC, ancestral population size), a demographic expansion occurred (NCC, current cultivated population size), TEXP (time expansion) generations ago. The wild population size is assumed to be constant through time (NWILD). The migration rate from wild to domesticated populations was also modeled, being equal to MIGWC = NMWC/NWILD, where NMWC is the number of wild migrants and NWILD is the wild population size (Fig. S1). The domestication bottleneck was modeled only for the Cult-TMVB population. For Cult-SMOCC and Cult-SUR-CH the modeled bottlenecks correspond to the traditional varieties spreading and it was assumed that they occurred after the initial domestication bottleneck. We did not find evidence of gene flow for the Cult-OV and Cult-TMVB-Spain (see results), therefore we modeled the ancestral population size (NANC), the time when a bottleneck started (TBOT), the population size during the bottleneck (NBOT), the demographic expansion time (TEXP), and the current population size (NCUR; Fig. S1).

We ran 100,000 simulations with 100 independent replicates for each model. For the best-fit model for those scenarios with gene flow, the likelihoods of the best runs were compared, estimating the AIC weight. Then we performed 50 bootstraps for the best-fit model to obtain the mean and 95% confidence intervals for each parameter.

## Results

### Sampling and SNV calling

After mapping, SNV calling and filtering, 237 individuals of *P. coccineus* (89 wild, 131 cultivated and 17 ferals), 20 of *P. vulgaris* and 35 of *P. dumosus* were kept. The mean missing data was 1.22%, and the mean depth per site was 31.04x. The SNV data set contained 79,286 SNVs, of which 11,019 variants were found in non-genic regions (13.90%), 35,429 within introns (44.68%) and 32,838 within CDS (41.42%). Regarding the variants within CDS, 13,738 (41.84%) were predicted as synonymous mutations, 18,392 (56.01%) as nonsynonymous, 541 (1.65%) as frameshift and 248 as nonsense (0.75%).

### Defined populations

The 237 samples from these 24 geographic locations were grouped into 15 populations (Fig. 1a; Fig. S2), based on the phylogenetic tree constructed with FastTree, the genetic groups identified by Admixture (eight genetic groups, half of these correspond to wild samples and the other half to traditional varieties; Fig. S2a and S3), and the PCA results (Fig. S2b and S4). The tree topology is similar to the one constructed by Guerra-García et al. (2017). Cultivars formed a monophyletic clade, and wild populations from the TMVB were the closest to the domesticated group.

**Fig. 1.**
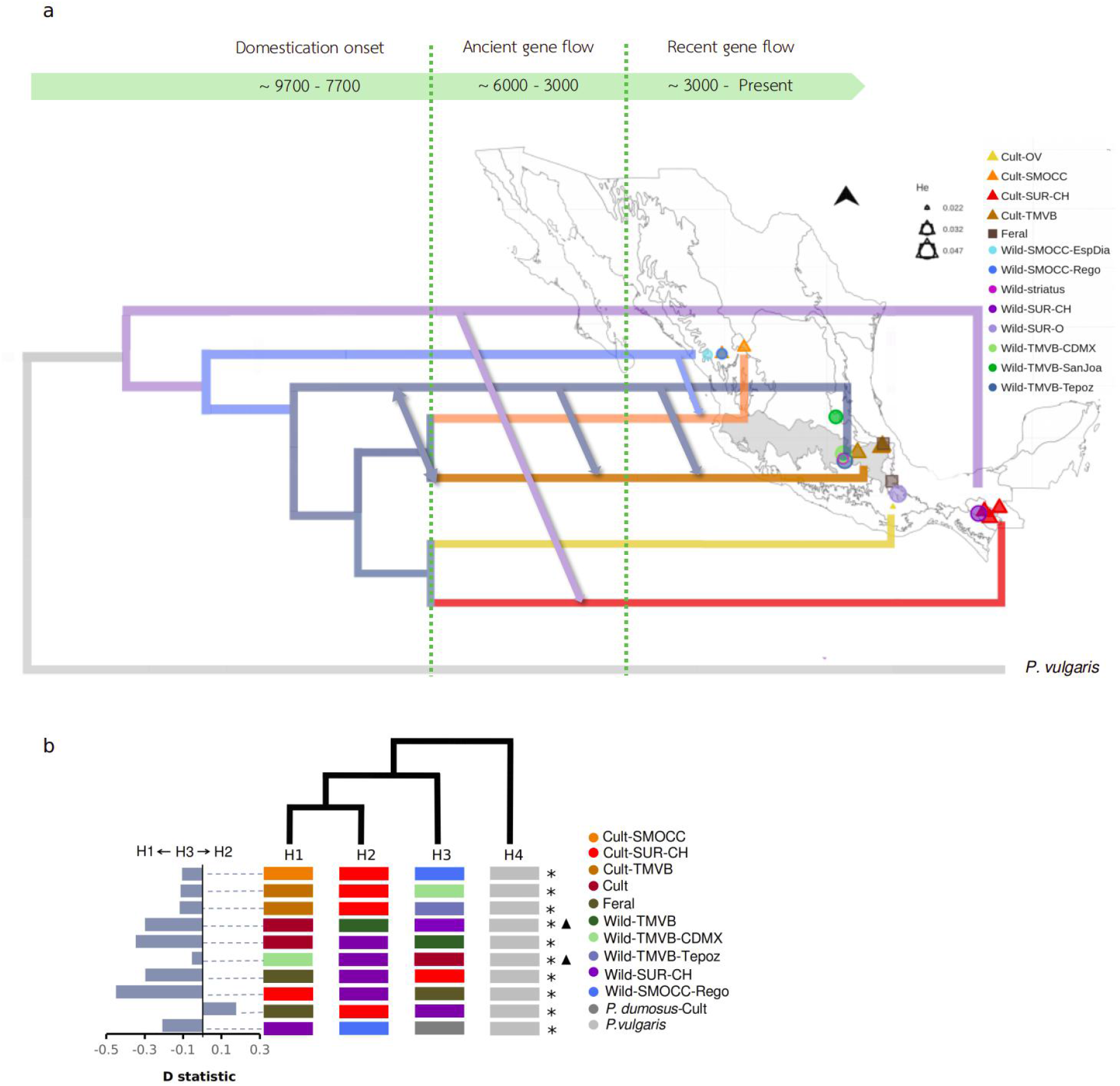
Population history of wild populations and traditional varieties of scarlet runner bean. a) Distribution map of genotyped populations in Mexico. Circles indicate wild populations and triangles show traditional varieties. Boundaries represent 21 ecoregions as defined by CONABIO (2008). The tree shows the phylogenetic relationship between the populations, arrows points indicate the direction of gene flow over time b) Gene flow test (ABBA-BABA). The tree shows the phylogenetic relations assumed for the gene flow scenarios. Asterisks indicate statistical significance after Bonferroni correction (*p* < 0.05), and black triangles point out the gene flow scenarios obtained with TreeMix. The plot on the left shows Patterson’s D-statistic: when D is positive, there is gene flow between H3 and H2; when D is negative, there is gene flow between H3 and H1.

The 15 populations comprised four ecoregions of Mexico (as defined in CONABIO 2008). Eight of these populations were made up of by wild individuals: two from Sierra Madre del Sur and Chiapas Highlands (Oaxaca Wild-SUR-O; and Chiapas Wild-SUR-CH); three from Trans-Mexican Volcanic Belt (Mexico City, Wild-TMVB-CDMX; San Joaquín, Wild-TMVB-SanJoa; Tepoztlán, Wild-TMVB-Tepoz); one identified as subsp. *striatus* (Wild-striatus); and two from the Sierra Madre Occidental (Regocijo, Wild-SMOCC-Rego; Espinazo del Diablo, Wild-SMOCC-EspDia). The other six populations corresponded to cultivars: from Sierra Madre del Sur (Cult-SUR-CH); Oaxaca Valley (Cult-OV); Trans-Mexican Volcanic Belt (Cult-TMVB); Sierra Madre Occidental (Cult-SMOCC); the Spain cultivar, which was grouped within traditional varieties from TMVB in the ancestry analysis (Fig. S2a; Cult-TMVB-Spain); and the breeding line Blanco-Tlaxcala, with ancestry from the SMOCC cultivars (Fig. S2a; Cult-SMOCC-BlaTla). Finally, all individuals identified as ferals were assigned to one group (Feral). The first word of the population name corresponds to the type of samples, followed by the genetic group assigned by Admixture, and the last letters indicate the population.

### Gene flow and wild-crop introgression

Three gene flow events were proposed by TreeMix (Fig. S5): 1) from an ancestral cultivar lineage to TMVB wild population; 2) an ancient event from Wild-SUR-CH to an old clade that included all cultivars; 3) from Wild-SUR-CH to ferals. These three scenarios were tested using the ABBA-BABA method and only the first two were supported (Fig. 1b).

We also looked for introgression among sympatric populations, and we found evidence of gene flow from wild populations to cultivars: from Wild-SMOCC-Rego to Cult-SMOCC; from Wild-TMVB-CDMX and Wild-TMVB-Tepoz to Cult-TMVB; from Wild-SUR-CH to Cult-SUR-CH (Fig. 1B). The only signal of introgression from cultivars to wild populations was the first scenario proposed by TreeMix (from ancestral cultivars to Wild-TMVB). In regard to the feral group, bidirectional gene flow among Feral and Cult-SUR-CH populations.

Sympatric populations of *P. dumosus* and *P. coccineus* occur in the Southern region of Mexico. Therefore, we tested for introgression between these two species. ABBA-BABA test supported introgression from *P. dumosus* to Wild-SUR-CH but not in the opposite direction, not even with the traditional varieties of the same region. Finally, the test showed evidence of frequent gene flow among traditional varieties (Table S2).

### Looking for domestication bottlenecks and demographic histories

We constructed an SFS for each population and SNV category. Patterns varied among populations, suggesting that they have gone through different evolutionary processes (Fig. S6). Most of the wild populations presented a slight excess of low-frequency alleles and subspecies *striatus* was the only wild population that showed a lack of low-frequency variants (Fig. S6). Three traditional varieties and ferals also presented an excess of low-frequency alleles (Cult-SMOCC, Cult-SUR, and Cult-TMVB), while Cult-TMVB-Spain showed a deficit. Nonsynonymous mutations were the most abundant variants at low-frequency in all populations (Fig. S6).

Higher ROH was found in cultivated populations compared to the wild ones (Fig. S7). Particularly the European cultivar (Cult-TMVB-Spain) had the largest ROH length, followed by the traditional variety from Oaxaca Valley (Cult-OV). This suggests that cultivated populations have gone through demographic bottlenecks, but the excess of low-frequency variants observed in several traditional varieties shows evidence of demographic expansions. Based on these results an initial bottleneck, followed by a demographic expansion were modeled for the cultivated populations using fastsimcoal. The gene flow found from wild to cultivars from SMOCC, TMVB and SUR-CH were integrated into the demographic models, testing introgression at different times (ancient, recent, and constant; Fig. 1a and Fig S1).

The best-fit scenario for Cult-TMVB includes constant introgression from the wild relatives to traditional varieties of this region, a severe bottleneck (NAC = ~2,500) associated with domestication time around 9,700 generations ago, followed by a relatively recent expansion (TEXP = ~1,500), and a current population size of ~759,000 (Fig. 1a, Fig. 2a and Table S2). The best-fit scenario for SMOCC populations suggests a recent introgression (from 3,000 generations ago to present), a less severe bottleneck associated with cultivar spreading (NAC = ~18,000) and a current population size of ~395,000. For the SUR-CH the best model predicted an ancient introgression (from 6,000 to 3,000 generations ago), a spreading bottleneck that led to an ancestral population size of ~8,500 and a current population of ~742,000 (Fig. 1a, Fig. 2b and Table S2). The SUR-CH population had the highest migration rate (MIGWC = 1.93E-05) of the three populations (Table S2).

**Fig. 2.**
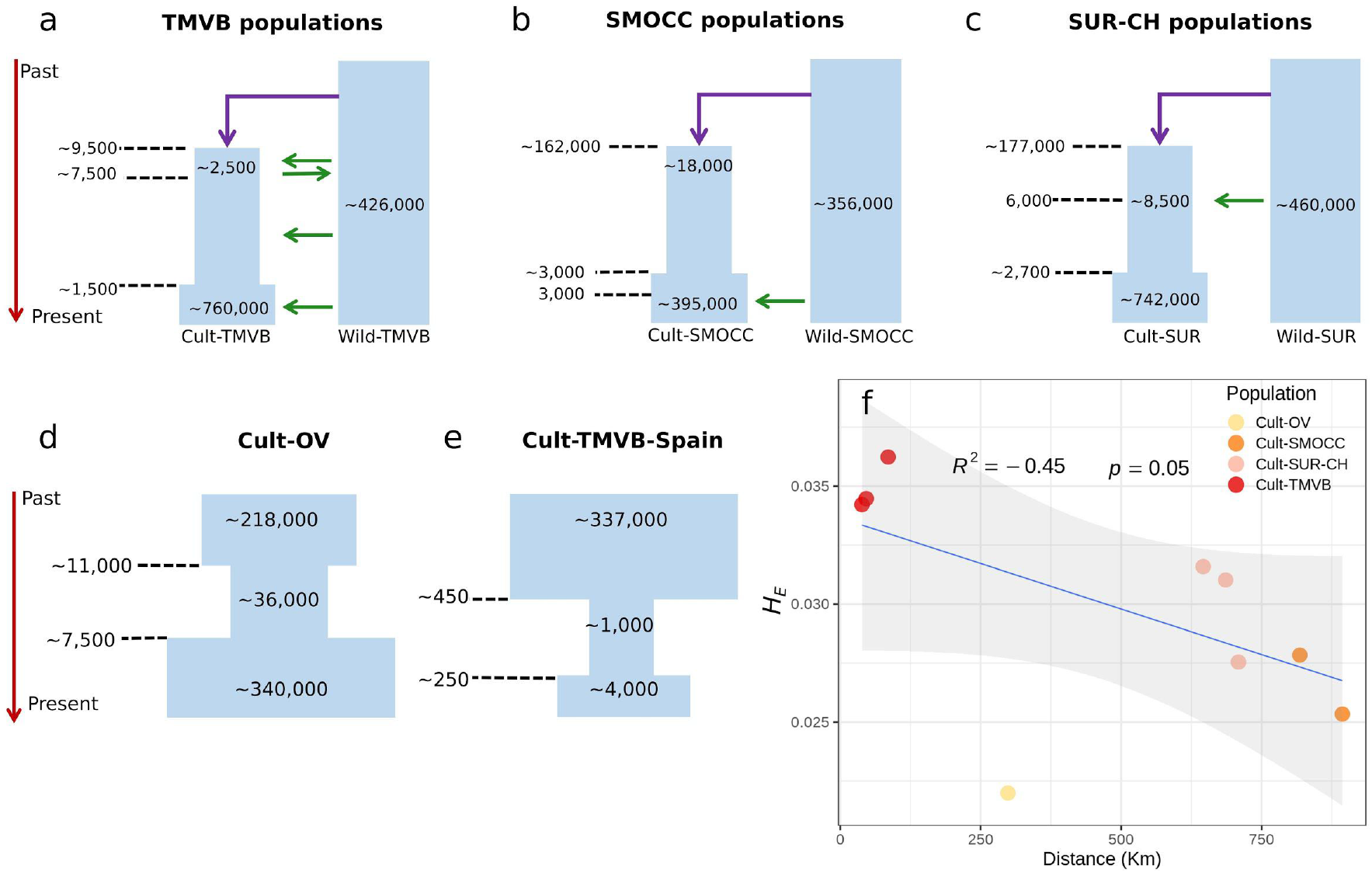
Best demographic scenarios and their parameters estimated with FastSimCoal. a) The best scenario for TMBV populations involves constant introgression; b) recent introgression in the SMOCC populations; and c) ancestral introgression in the case of populations from SUR-CH. For all populations the direction of the gene flow was from wild to cultivated beans. Ancestral bidirectional gene flow was included for the first 2,000 generations after the beginning of domestication only for TMVB, where domestication took place. Demographic models for Cult-OV (d) and Cult-TMVB-Spain (e), which have gone through bottlenecks in the absence of gene flow with our samples. f) Correlation between the genetic diversity (*H_E_*) and the distance from the centroid of the Cult-TMVB locations to traditional variety locations.

In the case of Cult-OV and Cult-TMVB-Spain only the severity and timing of the bottleneck were modeled. The bottleneck in Cult-OV was not as stringent as in all the other populations (NBOT = ~36,000), and its current population size is ~340,000 (Fig. 2d and Table S2). In contrast, the bottleneck that Cult-TMVB-Spain went through was severe (NBOT = ~1,000) and the current population size remains relatively low (NCC = ~4,000; Fig. 2e and Table S2).

### Measuring diversity and private alleles

Most wild populations showed the highest diversity. Both Wild-SUR populations (Oaxaca *H_E_* = 0.047; Chiapas *H_E_* = 0.044) and Wild-TMVB-CDMX (*H_E_* = 0.045) had the greatest amount of genetic diversity. Nevertheless, the Cult-TMVB and Feral populations showed high levels of variation, even higher than some wild populations (Wild-SMOCC and Wild-striatus). Cultivars from Oaxaca, Spain and the breeding line Blanco Tlaxcala showed the lowest genetic diversity. All cultivars showed a higher inbreeding coefficient (*F*_IS_) than wild populations (Figure S8).

When only segregating sites within each population were considered, a large proportion of private alleles was observed (Fig. S9). Private alleles were classified as: private to wild populations, to cultivars, to ferals, and to each population. The last class of private alleles was the most abundant. The proportion of private alleles was greater in wild samples. A full 54% of the segregating sites within Wild-SMOCC-EspDia and within Wild-SUR-O were private to those populations. Nevertheless, all cultivars presented this type of allele. The lowest proportions were found in Cult-TMVB-Spain and Cult-SMOCC-BlaTla (9% in both populations). In CDS regions the proportion of private alleles was 64% in the Wild-SMOCC-EspDia population and 54% in Wild-SUR-O.

Regarding the nonsynonymous/synonymous ratio, it was lower in the shared variants compred to the private ones, and it was particularly high in the cultivars (Table S3). The cultivars with the lowest nonsynonymous/synonymous ratio were Cult-TMVB-Spain and the breeding line Blanco Tlaxcala (Cult-SMOCC-BlaTla). Rarefaction analyses showed greater allelic richness and private allelic richness in wild populations (Fig. S10). Mean allelic richness per site was particularly high in Wild-TMVB-CDMX, Wild-TMVB-Tepoz and Wild-SUR-O but was also elevated in Feral and Cult-TMVB populations. On the contrary, Cult-TMVB-Spain showed the lowest allelic richness.

Finally, a negative correlation (*R*^2^ = −0.4, *p* = 0.05) was found between heterozygosity in traditional varieties and the distance to the centroid of the area where TMVB traditional varieties were collected (Fig. 2f).

## Discussion

### Demographic histories of scarlet runner bean populations

Genome-wide comparisons between wild and cultivars have been studied in several crops like maize (Hufford *et al*., 2012), rice (He *et al*., 2011; Huang *et al*., 2012), soybean (Lam *et al*., 2010; Li *et al*., 2013; Zhou *et al*., 2015), common bean (Schmutz *et al*., 2014; Bellucci *et al*., 2014; Vlasova *et al*., 2016; Rendón-Anaya *et al*., 2017), cucumber (Qi *et al*., 2013; Wang *et al*., 2018), and wheat (Haudry *et al*., 2007; Cavanagh *et al*., 2013). These have shown that the severity of bottlenecks vary substantially among species and, within species, even between gene pools (Bitocchi *et al*., 2013).

We made inferences about the demographic history of the scarlet runner bean and found evidence of the bottlenecks associated with the domestication and subsequent spread from its center of origin. Each population presents a unique history, with different severity and timing of bottleneck and demographic expansion. Moreover, we found that introgression from the wild relatives into cultivars is frequent, and it has occurred at a different rate and time across the populations included in this work.

Despite the relatively high genetic diversity found in the Cult-TMVB, the best demographic model suggests a strong bottleneck related to domestication. But the constant introgression from the wild populations might have contributed increasing levels of genetic variation. Furthermore, the Cult-TMVB population has recovered and has significantly increased its size, allowing the accumulation of new mutations. The estimated domestication time is reasonable (9,700 generations ago), even though it is higher than for common bean (~8,000 years ago; Gepts, 1998; Kaplan & Lynch, 1999).

On the other hand, our results suggest that the introgression from wild relatives has only taken place during the last 3,000 generations in the sympatric SMOCC populations. In contrast, the introgression in the SUR-CH seems to be older (6,000 - 3,000 generations ago; Fig. 1) and at a higher migration rate (Table S3). Furthermore, the Cult-SUR-CH current population size is almost the same as Cult-TMVB, indicating a conspicuous expansion. Contrasting, the Cult-OV population, where no evidence of gene flow was found, presented the less severe bottleneck and the lowest genetic diversity among Mexican traditional varieties. This finding suggests that introgression has incorporated and increased the genetic diversity in domesticated populations.

The estimated time when the bottleneck occurred in Cult-TMVB-Spain is within the expected time, after the foundation of the Viceroyalty of New Spain in 1525. During this time, an intense bidirectional exchange between Spain and what is now Mexico existed. The introduction of scarlet runner bean to Europe resulted in a relatively recent and intense bottleneck and, even though there is an increase in the population size, it is still low compared with Mexican traditional varieties and has the lowest *H_E_*. Nevertheless, because just one European cultivar was analyzed, no general pattern can yet be inferred about scarlet runner bean in Europe.

### Frequent and asymmetric introgression from wild relatives to traditional varieties

Gene flow and introgression are frequent among *P. coccineus* populations, which may be facilitated by the sympatry of wild and domesticated populations.

Furthermore, our results suggest that it is asymmetric, being more common from wild to traditional varieties than in the opposite direction. Just one gene flow event from cultivar to wild population was found: from an old domesticated clade to Wild-TMVB (Fig. 1b and Fig. S5).

Crop dispersion from its domestication center implies adaptation to new environments, and because wild populations are presumably adapted to local conditions, introgression from wild to traditional varieties could be a source of adaptive variation (Janzen *et al*., 2019). Adaptive introgression has been reported for: maize, based on both morphological (Wilkes, 1977) and molecular evidence (Hufford et al., 2013; van Heerwaarden et al., 2011); barley (Poets *et al*., 2015); cassava (Bredeson *et al*., 2016); and potato, where in certain cultivars wild ancestry was estimated upward of 30% (Hardigan *et al*., 2017). Thus, adaptive introgression may explain the patterns of asymmetric gene flow observed in *P. coccineus* and might maintain the relatively high genetic diversity found in scarlet runner bean traditional varieties, reducing the bottleneck consequences due to domestication. Conversely, natural selection may prevent the increase of domesticated alleles in wild populations. This hypothesis needs to be tested by looking at the specific regions that are introgressed and their possible functions.

Since a recurrent goal in breeding programs is to introgress adaptive traits from wild relatives into cultivated (Warschefsky *et al*., 2014), these already introgressed traditional varieties, with three different wild pools, become a powerful resource for valuable agronomic traits dissection. It has been noticed that crop-wild introgressed populations contain a mixture of wild and crop alleles that can be valued as an *in situ* germplasm resource in comparison with nonintrogressed populations (Ellstrand 2018). Therefore these populations could be mined for crop improvement of the scarlet runner bean.

Contrastingly, symmetric gene flow from crop to wild has been reported in common bean (Papa & Gepts, 2003) and lima bean (Félix *et al*., 2014), resulting in the displacement and reduction of genetic diversity in the wild relatives. Papa *et al*. (2005) found a significantly higher differentiation between wild relatives and cultivars in parapatric and allopatric populations compared to sympatric ones. Furthermore, differentiation was higher in genes related to domestication, which presented a higher diversity in wild populations. This suggests that selection was preventing introgression from domesticated to wild forms at target loci even though in other regions introgression was large due to the lack of selection against domesticated maladapted genes in early generation hybrids between wild and cultivated individuals (Papa *et al*., 2005). The asymmetric gene flow in common bean was recently confirmed with genomic data (Rendón-Anaya *et al*., 2017). Conversely, the low introgression in the domesticated population is probably due to the identification of the F1 hybrids within a cultivated field by farmers with a mechanism similar to an incompatibility barrier (Papa *et al*., 2005). Indeed, a powerful mechanism that may be playing a role in the asymmetric gene flow is cross-incompatibility, which could be a side effect of domestication in scarlet runner bean. Cross-incompatibility is the interaction (or lack of it) between pollen and pistil that prevents the formation of hybrids, and could be unidirectional or bidirectional (de Nettancourt, 2001; Maune *et al*., 2018). An asymmetric cross-incompatibility system would prevent recombination and would be compatible with a genome-wide pattern. Indeed, asymmetric cross-incompatibility is acting between *P. coccineus* and *P. vulgaris* (Wall, 1970; Shii *et al*., 1982). It has been proposed that cross-incompatibility is a mechanism that avoids gene flow between maize varieties and its wild relative teosinte, and at least three incompatibility systems have been described (Kermicle *et al*., 1990; Evans & Kermicle, 2001; Kermicle & Evans, 2010; Padilla García *et al*., 2012).

More recently it has been shown that cross-incompatibility is not enough to isolate maize modern varieties, landraces and wild relatives (Padilla-García *et al*., 2016). Deeper knowledge about the mating system in wild *P. coccineus* and how it has changed due to domestication is needed to understand if cross-incompatibility occurs in scarlet runner bean.

#### Genetic diversity

As expected, the greatest values of genetic diversity were found in wild populations. Nevertheless, there are wild populations with lower genetic diversity than traditional varieties, such as both Wild-SMOCC and Wild-striatus.

The expectation that high genetic diversity will be maintained close to the center of domestication and decrease with increasing geographic distance is observed in our data (Fig. 2f). This supports the hypothesis proposed by Guerra-García *et al*. (2017) that domestication took place in the TMVB, which was the most diverse traditional variety. Conversely, populations that have gone through subsequent bottlenecks have had a shorter time to accumulate new variation. Additionally, it is likely that the cultivars from Spain (Cult-TMVB-Spain) and Blanco Tlaxcala breeding line (Cult-SMOCC-BlaTla) were under stronger artificial selection.

Populations with the highest genetic diversity in terms of *H_E_* present the highest allele richness, but not the greatest private allelic richness nor the highest proportion of private alleles. This was the case of Wild-TMVB-CDMX with the highest diversity values, but relatively low private alleles. In contrast, the proportion of segregating sites that were private to Wild-SMOCC-EspDia and Wild-SUR-O were remarkably high (46% for both groups). The presence of private alleles might be at least partially explained by population histories. An ancient population, with high population sizes and/or isolated, would show a high proportion of private alleles (Nielsen & Slatkin, 2013). This could be the case of the wild and highly diverse populations from Southern Mexico. But this is not the case of Wild-SMOCC-EspDia, and most of the private alleles are at low frequencies, suggesting that they originated recentlyin terms of the origin of the wild populations, which is likely pre domestication. The demography of the wild populations of scarlet runner bean remains elusive and a future demographic study of the wild relatives would complement and allow a deeper understanding of the genetic variation in *P. coccineus* as a species.

The proportion of private alleles is lower in traditional varieties compared to their wild relatives, but still significant. For example, in Cult-SUR and Cult-TMVB private variation represents 33% of their segregating sites. The demographic expansions of the traditional varieties that occurred after the domestication and spread bottlenecks have probably favored the emergence of new variants within the cultivars.

Several factors are acting together promoting and maintaining genetic diversity in scarlet runner bean, such as demographic expansions and introgression from the wild relatives. Furthermore, it has been suggested that domestication bottlenecks might be less severe for perennials than for annual plants because perennial species frequently have a cross-pollination mating system and overlapping generations (Gaut *et al*., 2015). Although scarlet runner bean is commonly cultivated as an annual crop, it is a perennial open-pollinated species, and occasionally its subterraneous structures are kept for the next agricultural cycle (Delgado-Salinas, 1988).

### Purifying selection acting in wild and domesticated populations

Most wild populations and traditional varieties presented an excess of low-frequency variants (Fig. S6). Furthermore, in all populations, the proportion of non-synonymous at low-frequency is higher than the SNVs within non-genic regions and synonymous mutations. This is the expected pattern under purifying selection avoiding the frequency increase of deleterious mutations (Nielsen & Slatkin, 2013). The nonsynonymous/synonymous ratio of the segregating sites is > 1 in all wild and cultivated populations, except in Cult-TMVB-Spain. Furthermore, this ratio is higher in the private segregating sites, which probably are the more recent variants in both wild and cultivated populations, noting that “recent” for both types of populations refer to different time intervals (Table S3). This might suggest the presence of genetic load both in wild populations and traditional varieties, mainly integrated by recent private variants. Negative selection is presumably acting by keeping these slightly deleterious variants at low frequencies within the populations.

The cultivars that showed the lowest nonsynonymous/synonymous ratio were Cult-TMVB-Spain (0.957), followed by the breeding line Blanco Tlaxcala (Cult-SMOCC-BlaTla, 1.037). The severe bottleneck that occurred during the introduction of scarlet runner bean to Europe probably increased inbreeding and the strong artificial selection during the development of the breeding line Blanco Tlaxcala possibly resulted in the expression and posterior purge of some deleterious alleles. González *et al*. (2014) reported inbreeding depression in European scarlet runner bean cultivars, which affected germination, survival rates, yield, and seed weight. This may indicate that although a genetic purge might have occurred, deleterious variants associated with complex or quantitative traits, like germination, yield and seed weight, were maintained. When inbreeding depression is caused by a small number of recessive alleles with major deleterious effects on fitness, rapid response to selection is expected. However, deleterious variants with small effects are less easily purged and can be maintained in the population (Byers & Waller, 1999; Charlesworth & Willis, 2009).

## Conclusions

The demography of crops shapes patterns and levels of genetic variation, on which natural and artificial selection can act. A deeper understanding of the demographic dynamic of crops not only helps us to make inferences about the domestication history but also allows us to properly explore and take advantage of the genetic diversity that is available.

Despite the high genetic variation found in most traditional varieties of scarlet runner bean, our results suggest that domestication led to a severe bottleneck, followed by a significant demographic expansion. Furthermore, the introgression of alleles from wild populations to the traditional varieties has played an important role in increasing genetic diversity. If wild relatives are adapted to local conditions, then potentially adaptive introgression might have played in the diversification of traditional varieties of *P. coccineus* and their range expansion after domestication. This hypothesis needs to be tested and the identification of the introgressed genes or genomic regions and their functions can provide evidence of adaptive introgression.

## Acknowledgments

We thank Myriam Campos, Erick García, Verónica González, Alfredo Villarruel, Nancy Gálvez and Rocío González for fieldwork assistance, Tania Garrido for laboratory technical assistance, Ernesto Campos Murillo and Alicia Mastretta for bioinformatic assistance and Alfonso Delgado for his guide and feedback. We acknowledge funding from the CONACYT grant number 247730, PAEP to AGG and IEUNAM to DP. Statistical analyses were carried out in the CONABIO’s computing cluster, which was partially funded by Secretaría de Medio Ambiente y Recursos Naturales (SEMARNAT) through the grant “Contribución de la Biodiversidad para el Cambio Climático” to CONABIO. Finally, we deeply thank all farmers for sharing with us their seeds and maintaining agrodiversity.

