## Supplementary material for "The genomic signature of wild-to-crop introgression during the domestication of scarlet runner bean (*Phaseolus coccineus* L.)"

**Table S1.** *Phaseolus coccineus* sampling material used. "Population" column refers to the 15 defined populations that were the clustering factor in the analysis. All samples corresponded to *P. coccineus* subsp. *coccineus*, except a wild population from Tres Marías, Morelos. The last column shows the number of individuals that were kept after variant filtering.

| State and location | Status | Population | Latitude | Longitude | Altitude | *n* |
|---|---|---|---|---|---|---|
| Oaxaca, Cuilapam | Cultivar | Cult-OV | 16.99 | -96.78 | 1577 | 13 |
| Durango | Cultivar | Cult-SMOCC | NA | NA | NA | 8 |
| Durango, Regocijo | Cultivar | Cult-SMOCC | 23.68 | -105.12 | 2566 | 5 |
| Durango, Villa Unión | Cultivar | Cult-SMOCC | 23.97 | -104.04 | 1901 | 12 |
| Blanco Tlaxcala | Breeding line | Cult-SMOCC-BlTl | NA | NA | NA | 12 |
| Chiapas, González León | Cultivar | Cult-SUR-CH | 16.51 | -92.06 | 1579 | 10 |
| Chiapas, Nahá | Cultivar | Cult-SUR-CH | 16.94 | -91.59 | 917 | 12 |
| Chiapas, Oxchuc | Cultivar | Cult-SUR-CH | 16.80 | -92.32 | 2001 | 12 |
| Veracruz, Frijol Colorado | Cultivar | Cult-TMVB | 19.59 | -97.35 | 2419 | 11 |
| Veracruz, Orilla del Monte | Cultivar | Cult-TMVB | 19.66 | -97.29 | 2402 | 10 |
| Puebla, Tlalancaneca | Cultivar | Cult-TMVB | 19.36 | -98.51 | 2372 | 12 |
| Puebla | Cultivar | Cult-TMVB | NA | NA | NA | 6 |
| Spain | Cultivar | Cult-TMVB-Spain | NA | NA | NA | 8 |
| Veracruz, Altotonga | Feral | Feral | 19.75 | -97.25 | 1959 | 9 |
| Oaxaca, Huautla | Feral | Feral | 18.10 | -96.83 | 1746 | 8 |
| Durango, Espinazo del Diablo | Wild | Wild-SMOCC-EspDia | 23.64 | -105.82 | 1422 | 11 |
| Durango, Regocijo | Wild | Wild-SMOCC-Rego | 23.68 | -105.12 | 2566 | 12 |
| Morelos, Tres Marías | Wild Subsp. *striatus* | Wild-striatus | 19.10 | -99.21 | 3024 | 16 |
| Chiapas, San Cristóbal | Wild | Wild-SUR-CH | 16.70 | -92.60 | 2229 | 7 |
| Oaxaca, Comaltepec | Wild | Wild-SUR-O | 17.55 | -96.53 | 2876 | 4 |
| Ciudad de México, REPSA | Wild | Wild-TMVB-CDMX | 19.32 | -99.20 | 2328 | 11 |
| Ciudad de México, Tlalpan | Wild | Wild-TMVB-CDMX | 19.29 | -99.19 | 2334 | 8 |
| Querétaro, San Joaquín | Wild | Wild-TMVB-SanJoa | 20.93 | -99.56 | 2381 | 6 |
| Morelos, Tepoztlán | Wild | Wild-TMVB-Tepoz | 19.00 | -99.13 | 1953 | 13 |

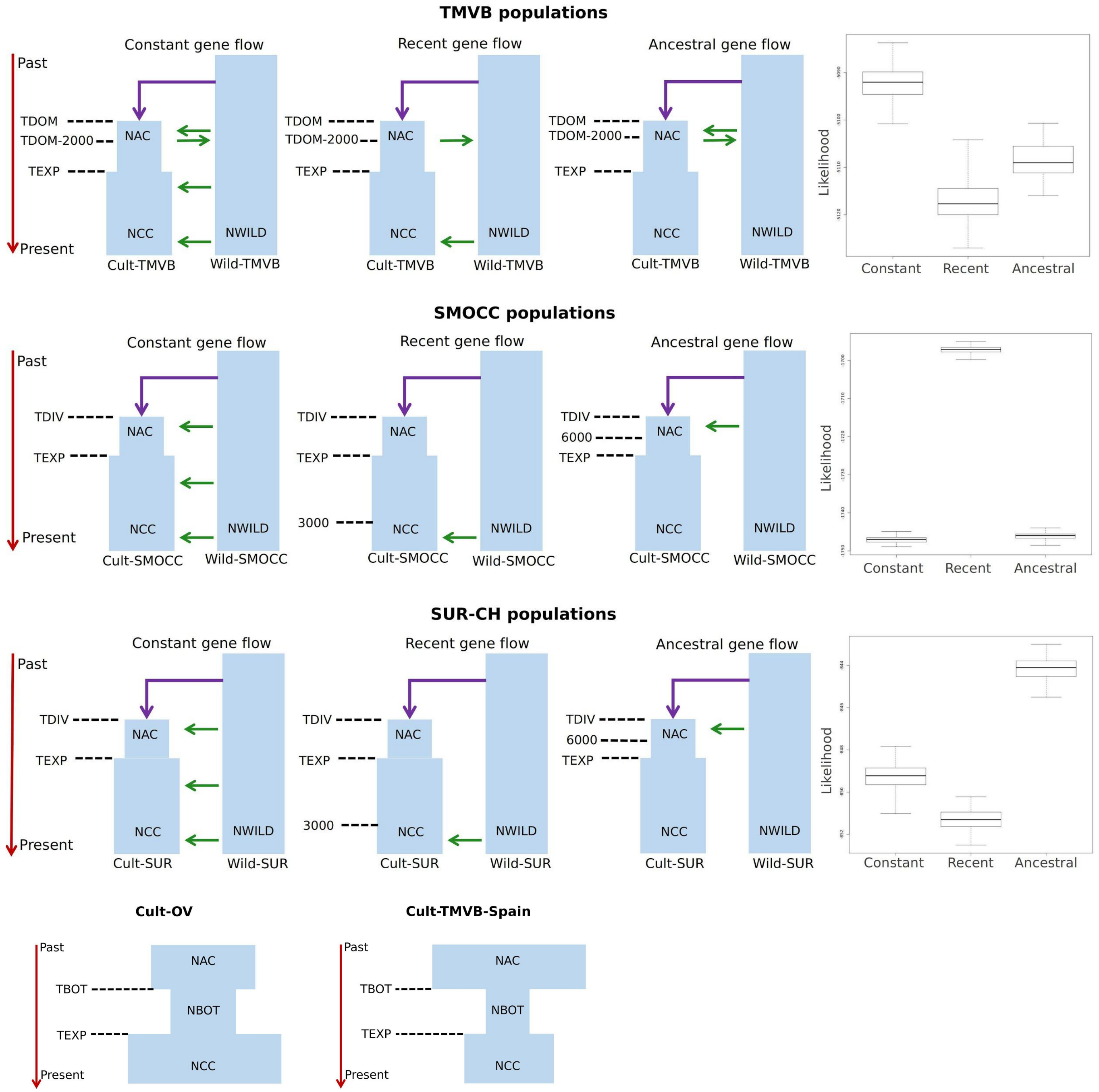


**Fig. S1.** Demographic models that were tested using FastSimCoal. For TMVB, SMOCC and SUR populations recent, constant and ancestral gene flow were tested. The likelihood for each of these scenarios are shown in the right column. For Cult-OV and the populations from Spain the severity and time of the bottleneck was estimated. NWILD= *Ne* wilds; NCC= Current *Ne* cultivars; NAC= Ancestral *Ne* cultivars; NBOT= *Ne* during the bottleneck; TBOT= bottleneck time (generations); TEXP= time of demographic expansion; TDOM= domestication time; TDIV= divergence time.

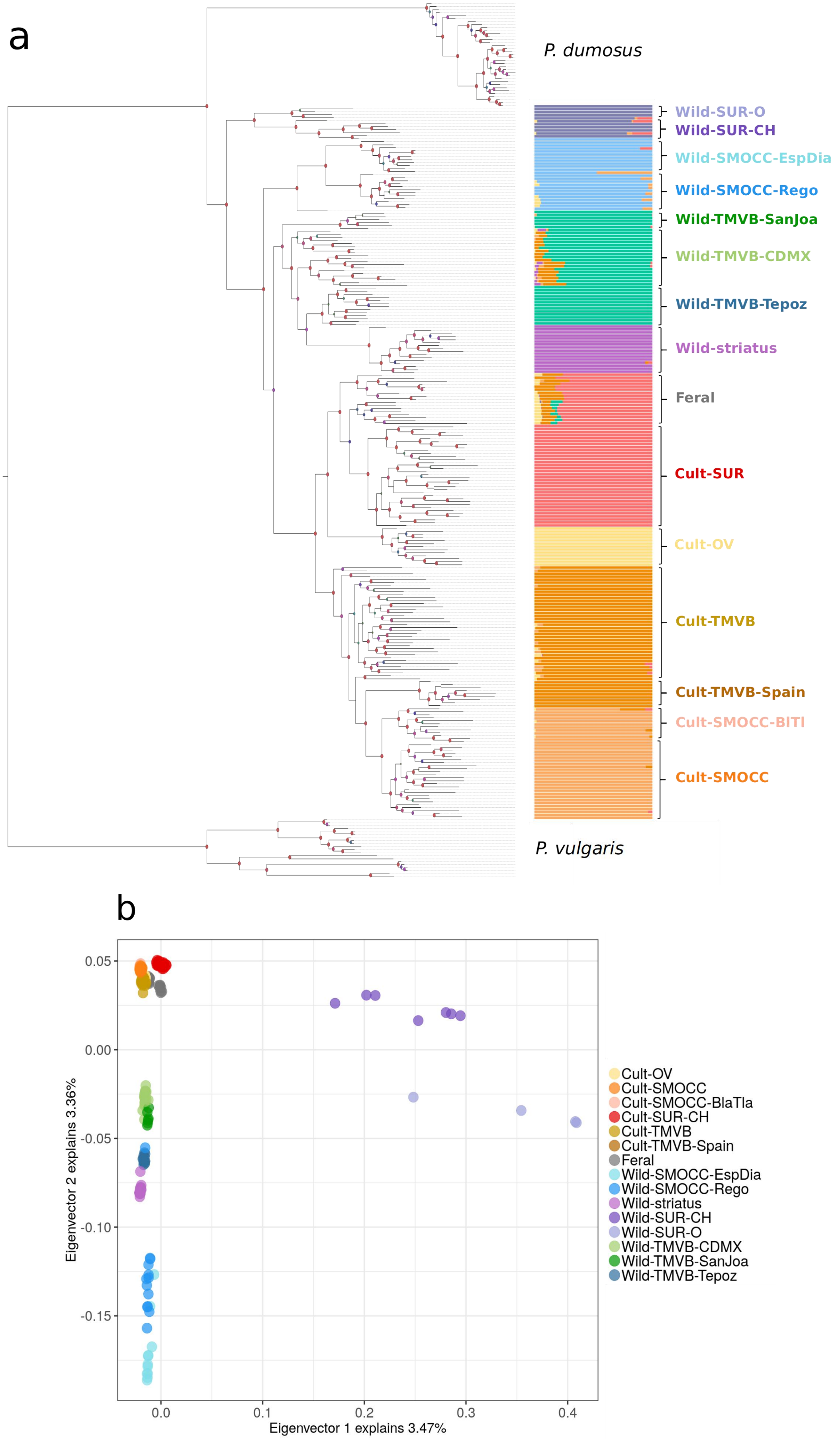


**Fig. S2.** a) Phylogenetic relationship among individuals of *P. coccineus* from Mexico and Admixture bar plot (shades of gray show the eight genetic groups) indicating the 15 defined populations. b) PCA plot for the first two principal components including all samples.

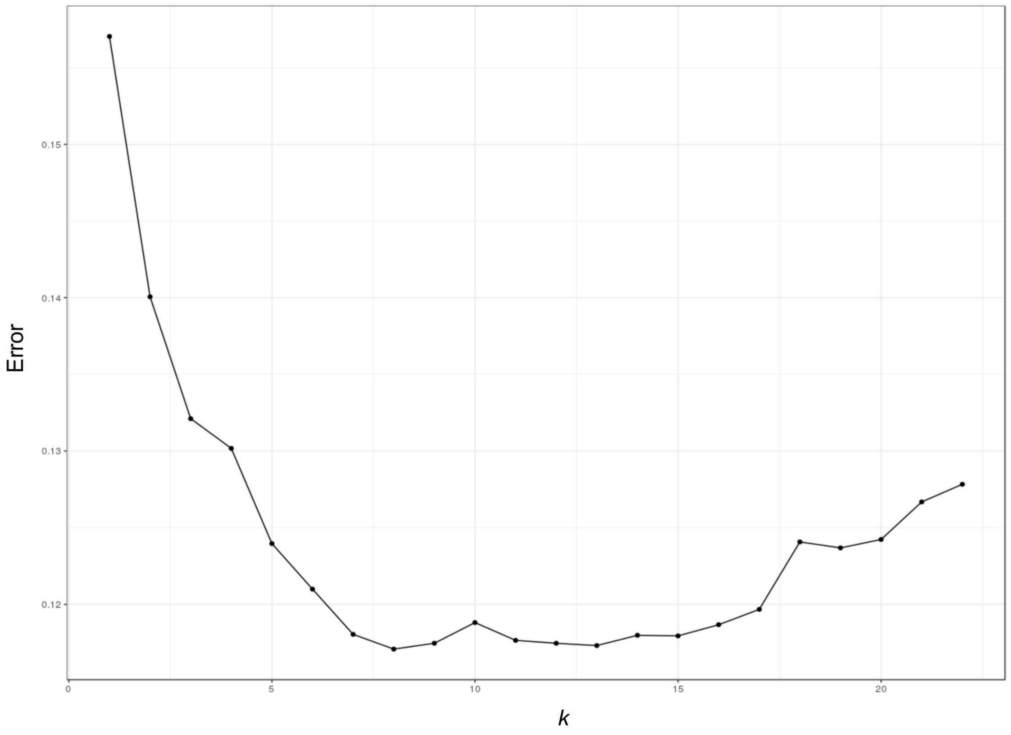


**Fig. S3.** Cross-validation errors among K values using Admixture software for the *P. coccineus* data set.

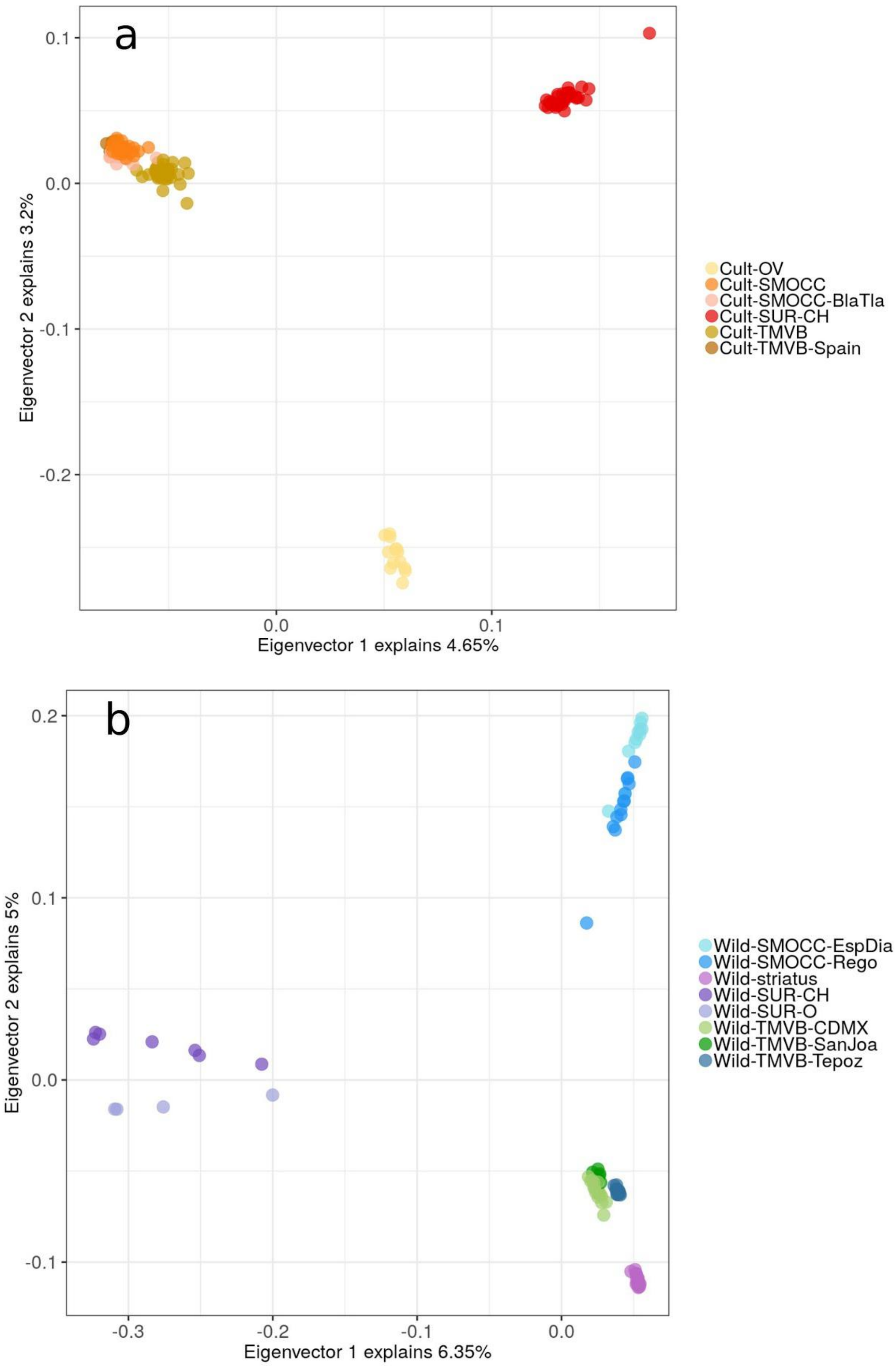


**Fig. S4.** PCA plots for the first two components of (A) cultivated and (B) wild populations.

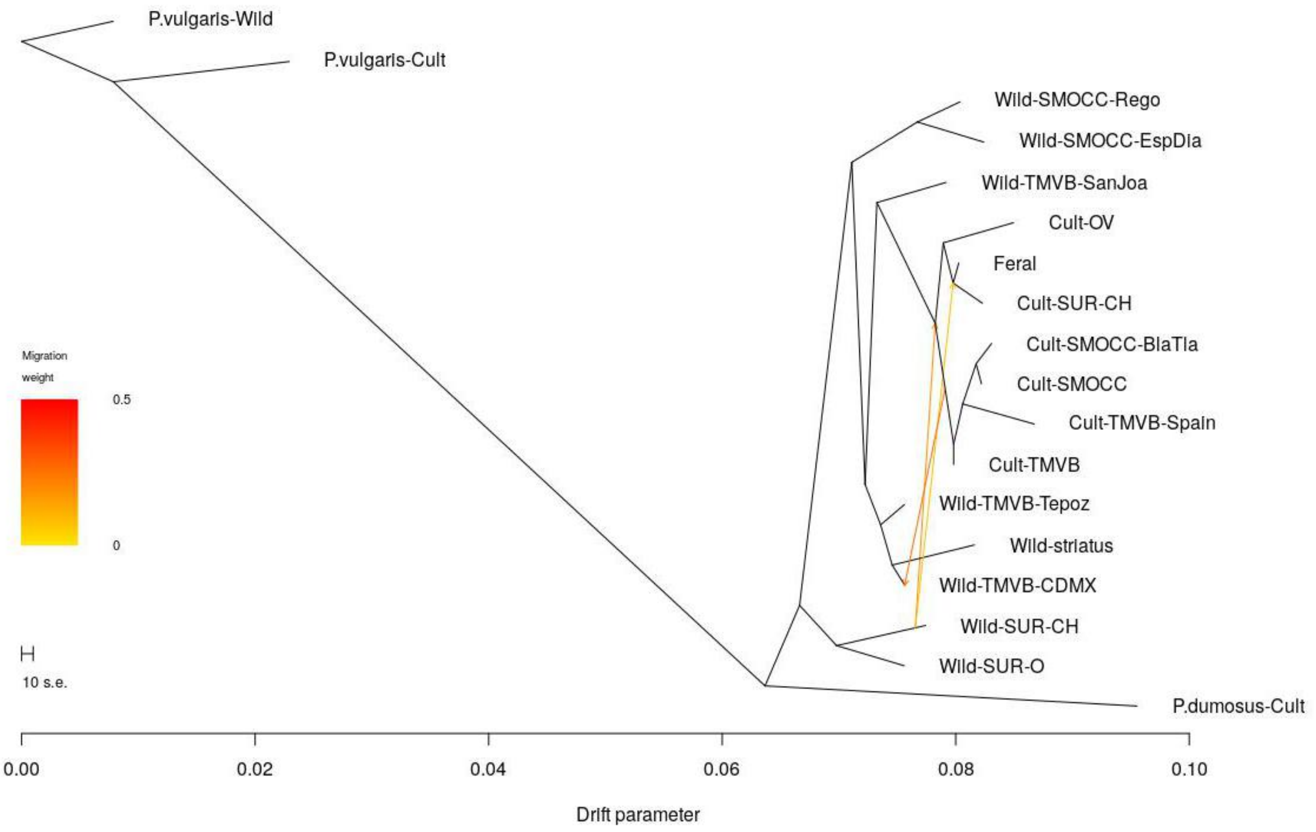


**Fig. S5.** Gene flow scenarios inferred by TreeMix. These scenarios were then tested with the ABBA-BABA approach.

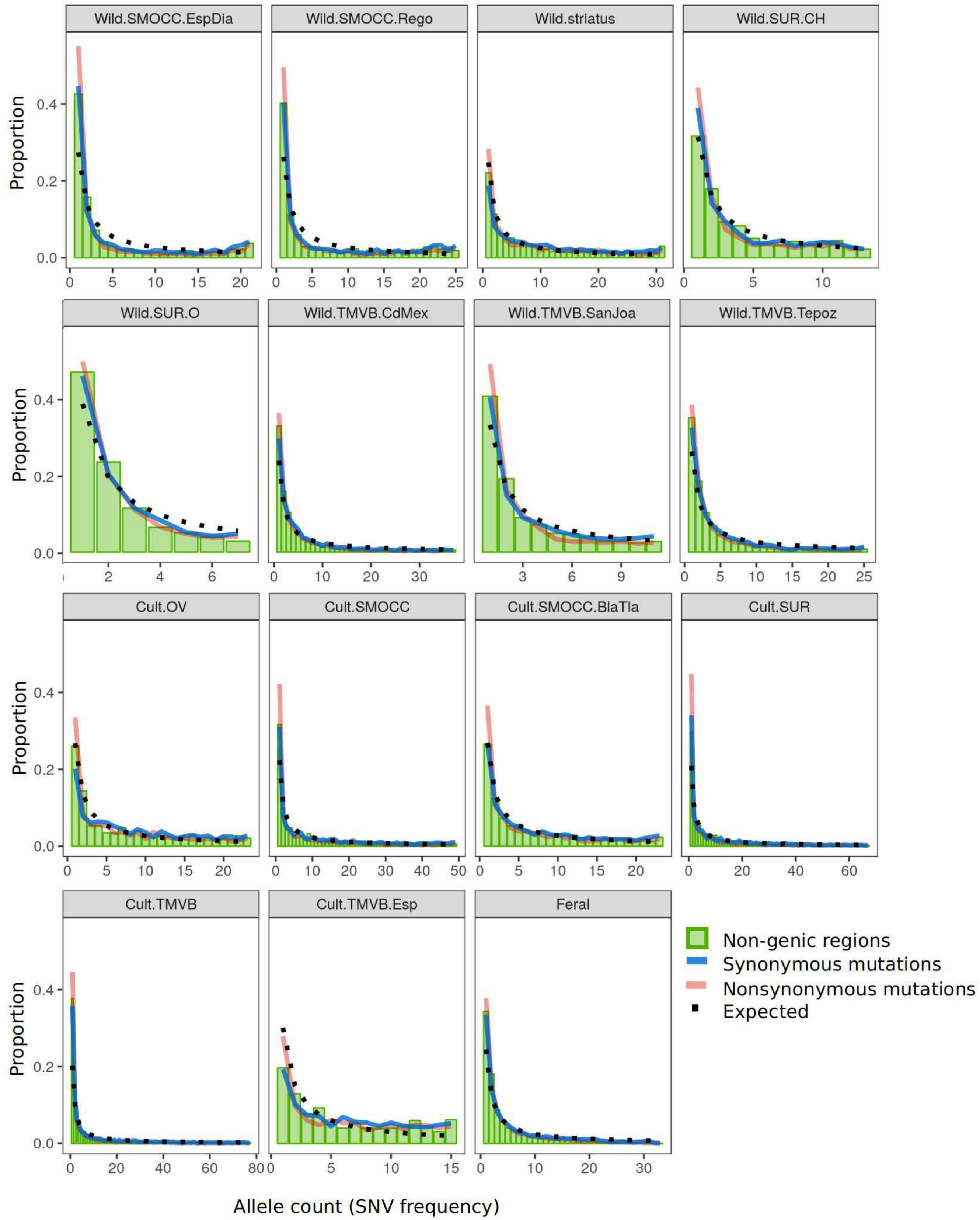


**Fig. S6**. Site Frequency Spectrum (SFS) calculated for each population. Colors indicate the SNV category. Points show the expected distribution.

**Table S2.** Gene flow models tested for cultivated, feral and wild populations of *P. coccineus*.

| H1 | H2 | H3 | H4 | D | Z | *p* value | Bonferroni *p* value | Statistically significant |
|---|---|---|---|---|---|---|---|---|
| Wild-TMVB | Wild-SUR-CH | Cult | P.vulgaris-Wild | -0.055 | -4.633 | 0.000004 | 0.000012 | * |
| Cult-SUR-CH | Wild-SUR-CH | Cult-OV | P.vulgaris-Wild | -0.459 | -32.937 | 0 | 0 | * |
| Cult-SUR-CH | Cult-TMVB | Cult-OV | P.vulgaris-Wild | -0.065 | -6.264 | 0 | 0 | * |
| Cult-SUR-CH | Cult-SMOCC | Cult-OV | P.vulgaris-Wild | -0.016 | -1.512 | 0.130576 | 0.391728 | Not |
| Cult-SUR-CH | Wild-SMOCC-Rego | Cult-SMOCC | P.vulgaris-Wild | -0.333 | -26.998 | 0 | 0 | * |
| Cult-SUR-CH | Cult-OV | Cult-SMOCC | P.vulgaris-Wild | 0.036 | 3.348 | 0.000815 | 0.002445 | * |
| Cult-TMVB-Spain | Cult-TMVB | Cult-SMOCC | P.vulgaris-Wild | -0.032 | -3.365 | 0.000767 | 0.002301 | * |
| Cult-OV | Wild-SUR-CH | Cult-SUR-CH | P.vulgaris-Wild | -0.334 | -20.696 | 0 | 0 | * |
| Cult-OV | Cult-TMVB | Cult-SUR-CH | P.vulgaris-Wild | -0.109 | -11.235 | 0 | 0 | * |
| Cult-OV | Cult-SMOCC | Cult-SUR-CH | P.vulgaris-Wild | -0.052 | -4.799 | 0.000002 | 0.000006 | * |
| Cult-SMOCC | Wild-SMOCC-Rego | Cult-SUR-CH | P.vulgaris-Wild | -0.424 | -41.54 | 0 | 0 | * |
| Cult-TMVB | Wild-TMVB-CDMX | Cult-SUR-CH | P.vulgaris-Wild | -0.303 | -32.568 | 0 | 0 | * |
| Cult-TMVB | Wild-TMVB-Tepoz | Cult-SUR-CH | P.vulgaris-Wild | -0.399 | -39.962 | 0 | 0 | * |
| Cult-TMVB&SMOCC | Wild-SUR-CH | Cult-SUR-CH | P.vulgaris-Wild | -0.274 | -17.266 | 0 | 0 | * |
| Feral | Wild-SUR-CH | Cult-SUR-CH | P.vulgaris-Wild | -0.297 | -19.972 | 0 | 0 | * |
| Feral | Cult-TMVB | Cult-SUR-CH | P.vulgaris-Wild | -0.064 | -6.545 | 0 | 0 | * |
| Cult-SMOCC | Cult-TMVB-Spain | Cult-TMVB | P.vulgaris-Wild | -0.021 | -2.238 | 0.025241 | 0.075723 | Not |
| Cult-SUR-CH | Wild-TMVB-CDMX | Cult-TMVB | P.vulgaris-Wild | -0.196 | -17.981 | 0 | 0 | * |
| Cult-SUR-CH | Wild-TMVB-Tepoz | Cult-TMVB | P.vulgaris-Wild | -0.295 | -24.989 | 0 | 0 | * |
| Cult-SUR-CH | Cult-OV | Cult-TMVB | P.vulgaris-Wild | 0.045 | 4.604 | 0.000004 | 0.000012 | * |
| Feral | Cult-SUR-CH | Cult-TMVB | P.vulgaris-Wild | 0.009 | 1.053 | 0.292386 | 0.877158 | Not |
| Cult-SUR-CH | Wild-SUR-CH | Cult-TMVB&SMOCC | P.vulgaris-Wild | -0.427 | -30.714 | 0 | 0 | * |
| Cult-SMOCC | Cult-TMVB | Cult-TMVB-Spain | P.vulgaris-Wild | -0.053 | -5.537 | 0 | 0 | * |
| Cult-SUR-CH | Wild-SUR-CH | Feral | P.vulgaris-Wild | -0.45 | -32.523 | 0 | 0 | * |
| Cult-SUR-CH | Cult-TMVB | Feral | P.vulgaris-Wild | -0.073 | -7.119 | 0 | 0 | * |
| Wild-SUR-CH | Wild-SMOCC-Rego | P.dumosus-Cult | P.vulgaris-Wild | -0.208 | -14.592 | 0 | 0 | * |
| Cult-SMOCC | Cult-SUR-CH | Wild-SMOCC-Rego | P.vulgaris-Wild | -0.105 | -10.908 | 0 | 0 | * |
| Wild-SUR-CH | P.dumosus-Cult | Wild-SMOCC-Rego | P.vulgaris-Wild | -0.363 | -28.417 | 0 | 0 | * |
| Cult | Wild-TMVB | Wild-SUR-CH | P.vulgaris-Wild | -0.298 | -34.042 | 0 | 0 | * |
| Cult-SUR-CH | Cult-OV | Wild-SUR-CH | P.vulgaris-Wild | -0.148 | -14.643 | 0 | 0 | * |
| Cult-TMVB&SMOCC | Cult-SUR-CH | Wild-SUR-CH | P.vulgaris-Wild | 0.173 | 17.836 | 0 | 0 | * |
| Feral | Cult-SUR-CH | Wild-SUR-CH | P.vulgaris-Wild | 0.176 | 20.397 | 0 | 0 | * |
| Wild-SMOCC-Rego | P.dumosus-Cult | Wild-SUR-CH | P.vulgaris-Wild | -0.168 | -10.96 | 0 | 0 | * |
| Cult | Wild-SUR-CH | Wild-TMVB | P.vulgaris-Wild | -0.347 | -30.331 | 0 | 0 | * |
| Cult-TMVB | Cult-SUR-CH | Wild-TMVB-CDMX | P.vulgaris-Wild | -0.113 | -13.239 | 0 | 0 | * |
| Cult-TMVB | Cult-SUR-CH | Wild-TMVB-Tepoz | P.vulgaris-Wild | -0.118 | -13.024 | 0 | 0 | * |

**Tables S3.** Demographic parameters estimated using FastSimCoal (95% CI). NWILD= *Ne* wilds; NCC= Current *Ne* cultivars; NAC= Ancestral *Ne* cultivars; NBOT= Ne during the bottleneck; TBOT= bottleneck time (generations); TEXP= time of demographic expansion; TDOM= domestication time; TDIV= divergence time; REXP= expansion rate; MIGWC= migration rate from wild to cult (NMWC/NWILD); MIGCW= migration rate from cult to wild (NMCW/NAC); NMWC= migrants from wild to cult; NMCW= migrants from cult to wild.

| Pop-model | NWILD | NCC | NAC | NBOT | TBOT | TEXP | TDOM | TDIV | MIGWC | MIGCW |
|---|---|---|---|---|---|---|---|---|---|---|
| Cult-TMVB_Wild-TMVB-constant | 425,777 (337,027 - 487,500) | 759,246 (716,234 - 834945) | 2,423 (1,999 - 3,216) | NA | NA | 1,635 (1,522 - 1,775) | 9,701 (7889 - 2509) | NA | 1.38E-05 (1.14E-05 - 1.64E-05) | 6.19E-06 (2.66E-06 - 1.73E-05) |
| Cult-SMOCC_Wild-SMOCC-recent | 355,636 (277,313 - 465,669) | 395,155 (273,532 - 622,906) | 17,574 (13,343 - 21,085) | NA | NA | 3,166 (3,006 - 3581) | NA | 162,039 (132,247 - 198,564) | 5.76E-06 (4.74E-06 - 7.07E-06) | NA |
| Cult-SUR_Wild-SUR-CH-old | 459,744 (394,880 - 519,337) | 741,887 (558,292 - 921,143) | 8,571 (6,277 - 12,023) | NA | NA | 2,706 (2,167 - 2965) | NA | 176,689 (154,796 - 210,670) | 1.93E-05 (1.70E-05 - 2.20E-05) | NA |
| Cult-TMVB-Spain | NA | 4,130 (1,976 - 6793) | 337,078 (241,184 - 495,681) | 1,047 (177 - 2,382) | 447 (156 - 753) | 247 (66 - 497) | NA | NA | NA | NA |
| Cult-OV | NA | 340,417 (325,384 - 355,451) | 218,142 (197,358 - 238,925) | 35,663 (33,873 - 37,453) | 11,116 (8,842 - 13,389) | 7,498 (6,217 - 8,780) | NA | NA | NA | NA |

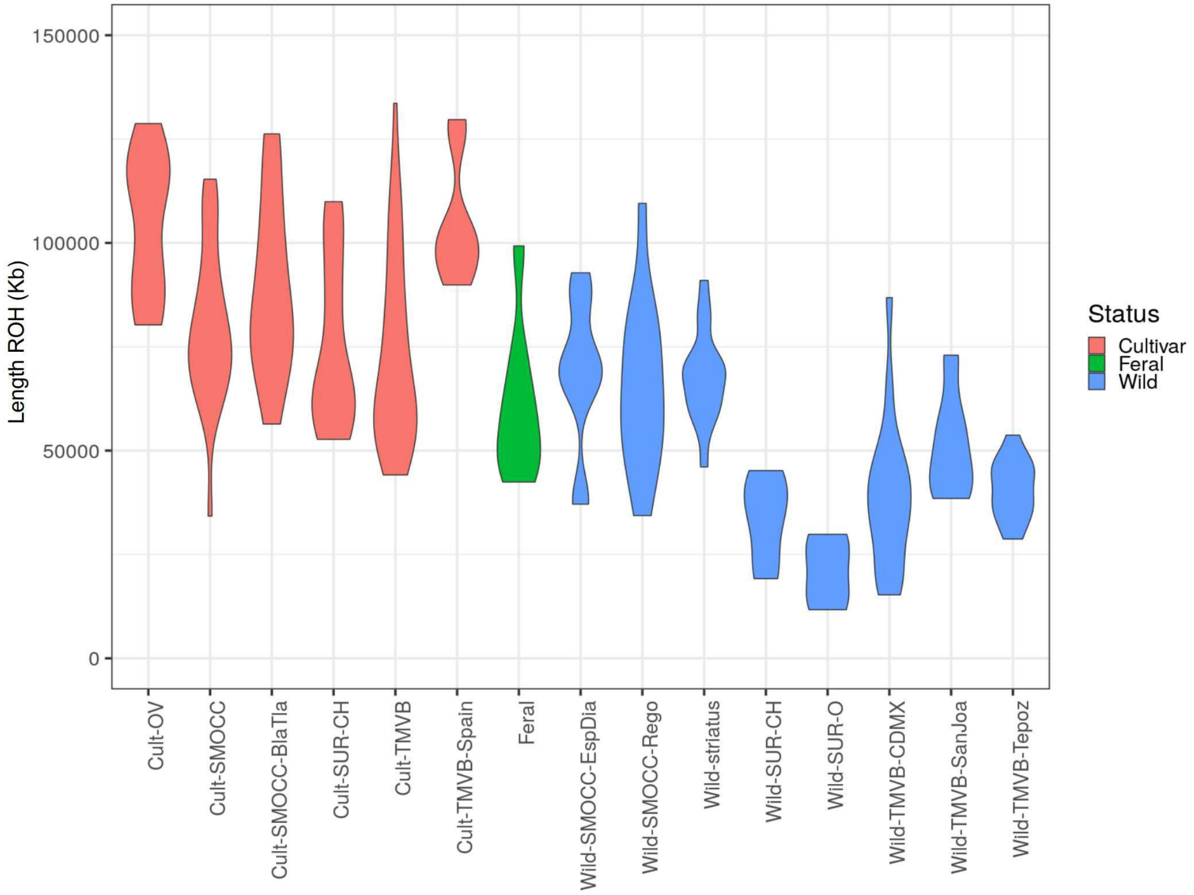


**Fig. S7.** Total length of ROH (Kb) estimated using a 500 Kb min window size. Colors indicate the type of *P. coccineus* sample.

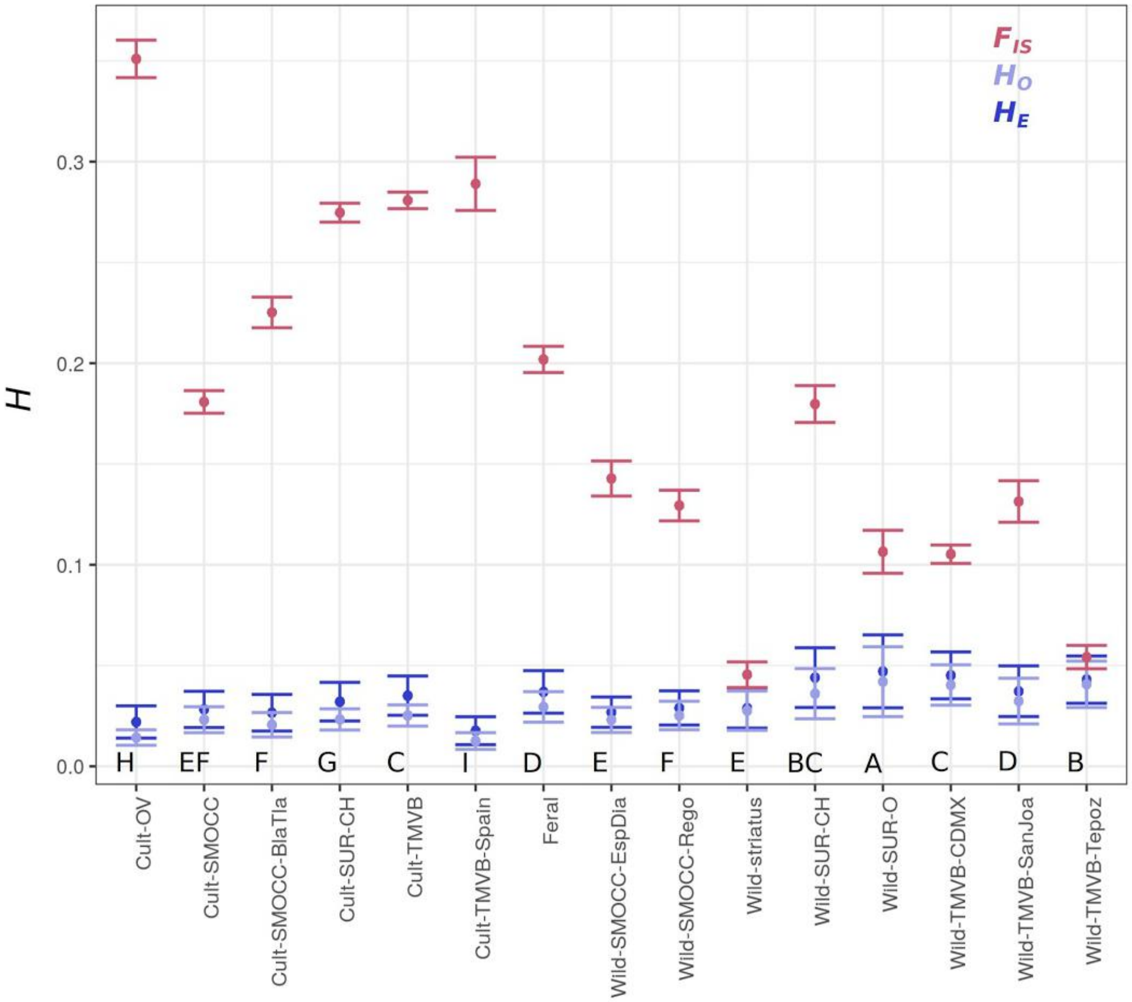


**Fig S8.** Genetic diversity (*H*) and inbreeding coefficient ($F_{IS}$; 95% CI) estimated for the 15 defined populations. Letters indicate populations that are statistically different in terms of $H_E$ and are shown in decreasing order.

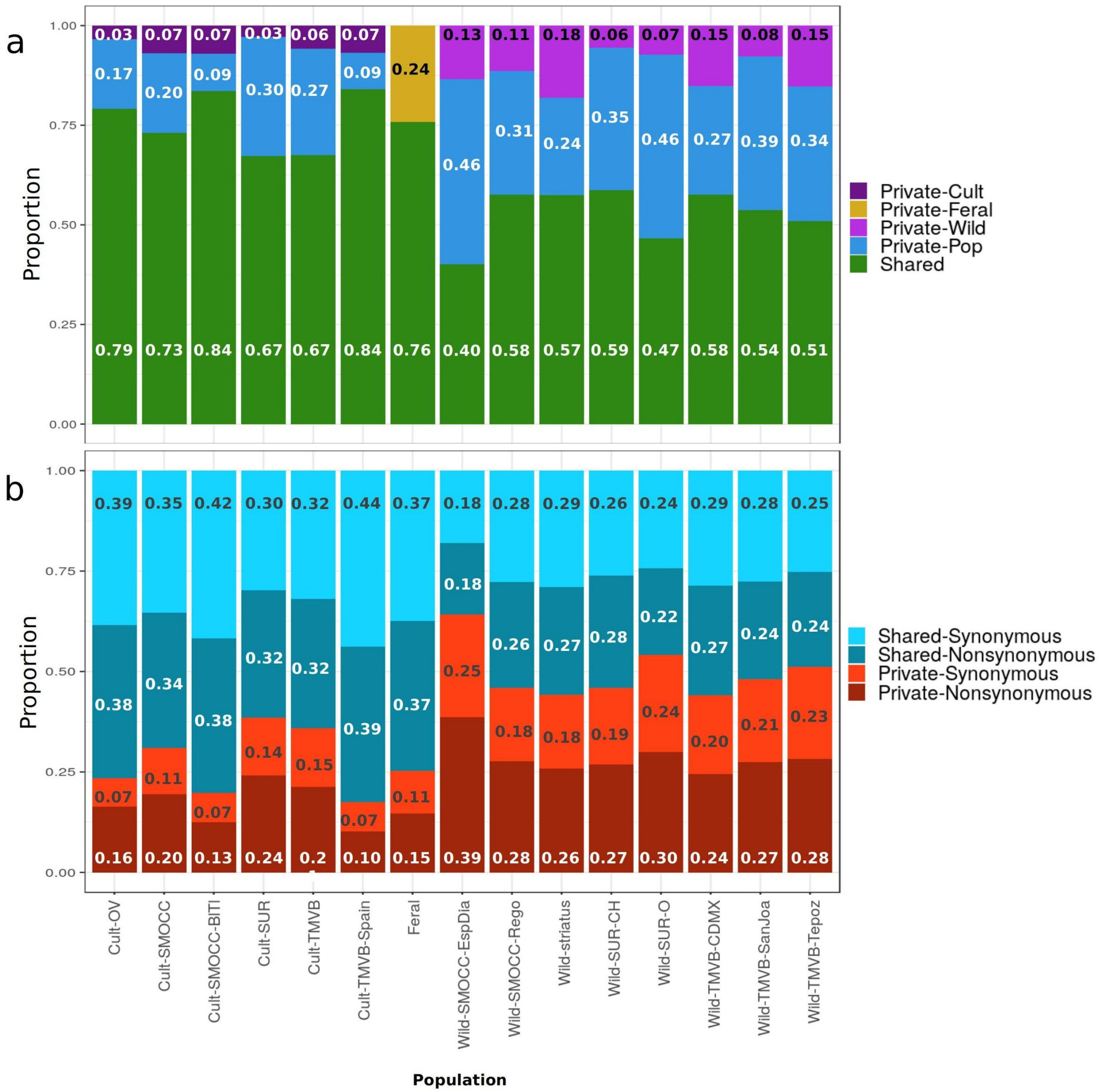


**Fig. S9.** A) Proportion of private and shares alleles for each population. B) Proportion of private and shared alleles within CDS regions, separating into synonymous and nonsynonymous mutations. The numbers inside the columns indicate the proportion of each category. Only segregating sites within populations are included.

**Table S2.** Proportion of segregating sites (SS) and nonsynonymous/synonymous ratio of SS splitted into the shared and private alleles found within each *P. coccineus* population.

| Population | Proportion SS | Nonsynonymous/synonymous | | |
|---|---|---|---|---|
| | | All SS | Shared SS | Private SS |
| Cult-OV | 0.080 | 1.196 | 0.991 | 2.306 |
| Cult-SMOCC | 0.140 | 1.138 | 0.953 | 1.708 |
| Cult-SMOCC-BlTl | 0.101 | 1.037 | 0.919 | 1.711 |
| Cult-SUR | 0.191 | 1.264 | 1.067 | 1.670 |
| Cult-TMVB | 0.237 | 1.148 | 1.008 | 1.452 |
| Cult-TMVB-Spain | 0.052 | 0.957 | 0.884 | 1.395 |
| Feral | 0.176 | 1.080 | 0.996 | 1.378 |
| Wild-SMOCC-EspDia | 0.134 | 1.294 | 0.986 | 1.511 |
| Wild-SMOCC-Rego | 0.138 | 1.174 | 0.948 | 1.519 |
| Wild-striatus | 0.102 | 1.113 | 0.928 | 1.405 |
| Wild-SUR-CH | 0.144 | 1.219 | 1.074 | 1.417 |
| Wild-SUR-O | 0.114 | 1.061 | 0.885 | 1.237 |
| Wild-TMVB-CDMX | 0.225 | 1.069 | 0.950 | 1.243 |
| Wild-TMVB-SanJoa | 0.117 | 1.067 | 0.876 | 1.320 |
| Wild-TMVB-Tepoz | 0.186 | 1.078 | 0.937 | 1.234 |

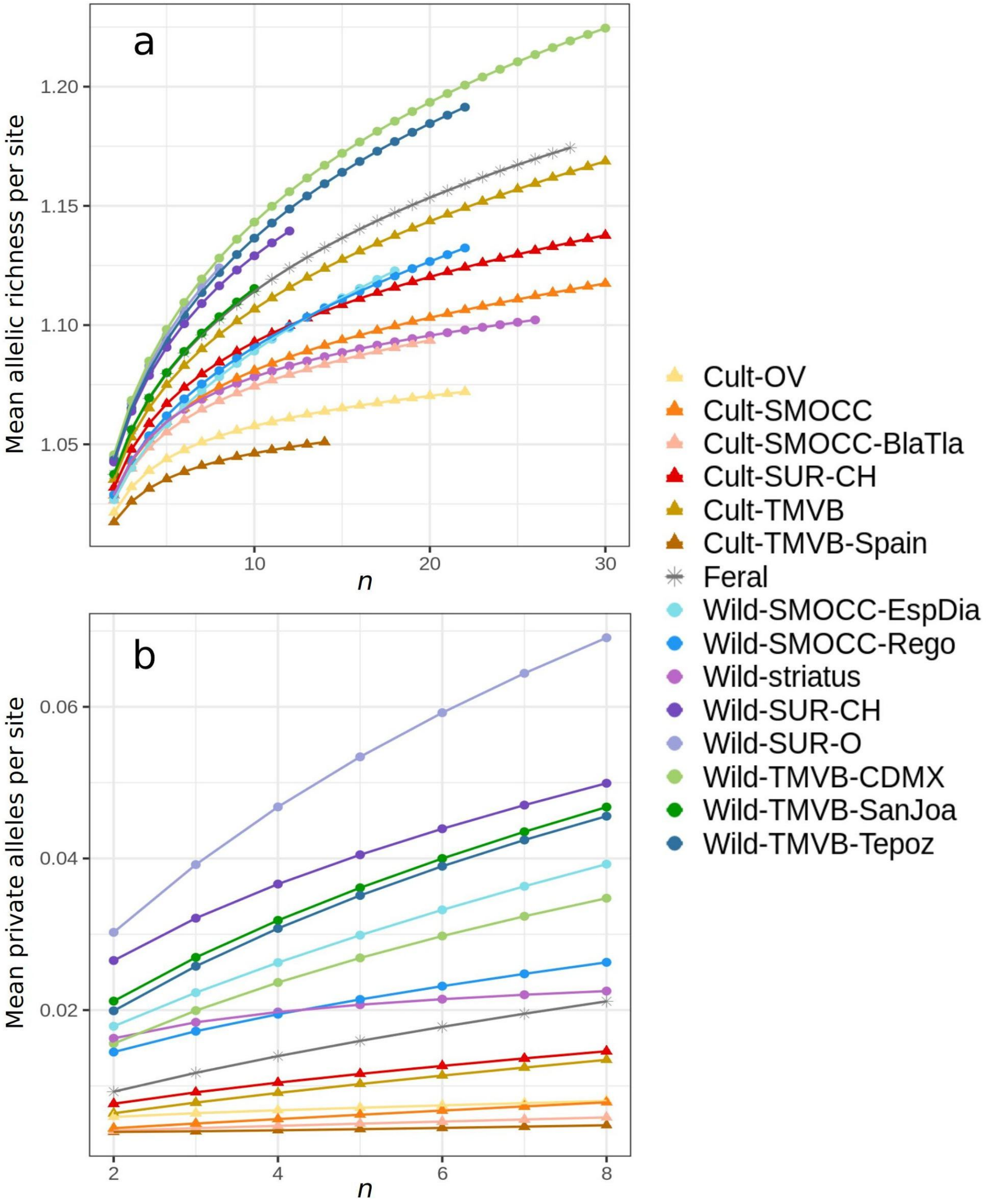


**Fig. S10.** A) Mean allelic richness per site and (B) mean private alleles per site estimated with AZDE for each population.
